# The impact of innate immunity on malaria parasite infection dynamics

**DOI:** 10.1101/2023.04.17.537202

**Authors:** Alejandra Herbert-Mainero, Philip J. Spence, Sarah E. Reece, Tsukushi Kamiya

## Abstract

Decades of research have investigated the molecular and cellular mechanisms that control the immune response to malaria. Yet, many studies offer conflicting results on the functional impact of innate immunity for controlling parasite replication early in infection. We conduct a meta-analysis to probe for consensus on the effect of innate immunity on parasite replication, examining three different species of rodent malaria parasite. Screening published studies that span four decades of research we collate, curate and statistically analyse infection dynamics in immune deficient or augmented mice to identify and quantify consensus and identify sources of disagreement among studies. Additionally, we estimate whether host factors or experimental methodology shape the impact of immune perturbations on parasite burden. First, we detected meta-analytic mean effect sizes (absolute Cohen’s h) for the difference in parasite burden between treatment and control groups ranging from 0.1498 to 0.2321 across parasite species. This range is considered a small effect size and translates to a modest change in parasitaemia of roughly 6-12% on average at the peak of infection. Second, we reveal that variation across studies using *P. chabaudi* or *P. yoelii* is best explained by stochasticity (due to small sample sizes) rather than by host factors or experimental design. Third, we find that for *P. berghei* the impact of immune perturbation is increased when young or female mice are used and is greatest when effector molecules (as opposed to upstream signalling molecules) are disrupted (up to an 18% difference in peak parasitaemia). Finally, we find little evidence of publication bias suggesting that our results are robust. The small effects sizes we observe, across three parasite species, following experimental perturbations of the innate immune system may be explained by redundancy in a complex biological system or by incomplete (or inappropriate) data reporting for meta-analysis. Alternatively, our findings might indicate a need to re-evaluate the efficiency with which innate immunity controls parasite replication early in infection. Testing these explanations is necessary to translate understanding from model systems to human malaria infections, manage immunopathology, and facilitate realism in mathematical models.

## 2 Introduction

The immune response plays a pivotal role in shaping the fate of infectious agents and host health outcomes (1). Innate immune effectors can block invasion or directly kill pathogens or the host cells that they reside within (e.g. via complement, phagocytosis or ROS). For naïve hosts that have no previous exposure to malaria parasites, the innate immune system triggers a rapid effector response that is thought to provide early control of pathogen replication by removing infected red blood cells (RBC) as well as short-lived extracellular parasites known as merozoites (2,3) This acute phase response is characterised by a pro-inflammatory environment with elevated concentrations of TNF and IFNγ, which can arrest parasite development and kill infected RBC, potentially buying time for the development of adaptive immune responses, which are essential for parasite clearance (4). However, establishing a causal link between specific innate effector mechanisms and host control of parasite replication *in vivo* has proved challenging (5) This is largely because innate effector molecules are pleiotropic (e.g. TNF and IFNγ can also suppress erythropoiesis to reduce red cell availability) (6,7)and because functional redundancy is a cardinal feature of the innate immune system. Consequently, it is difficult to find consensus on the extent to which innate immunity controls parasite replication and under what circumstances. To address this, we conducted a systematic and quantitative synthesis of the primary literature to ask how experimental manipulations of the innate immune system impact the dynamics of parasite replication in rodent models of malaria.

Following the discovery of rodent malaria parasites in the 1940s model systems have been developed to provide insight into all aspects of host-parasite interactions, including mechanistic knowledge of the immune response to infection (8,9)Rodent models offer a balanced compromise between manipulability and within-host ecological realism when compared to human infections and *in vitro* systems. Specifically, although studying human malaria offers the maximum clinical translational value, this is tempered by the difficulty of controlling for confounding factors in natural infections and an inability to manipulate the immune system *in vivo* (10). At the opposite end of the spectrum, *in vitro* model systems offer freedom of manipulation to probe molecular and cellular mechanisms in detail but lack realistic within-host processes that regulate dynamic immune response (11).Thus, despite important differences to human malaria rodent models can provide general insight into how host immunity controls acute infection (12).

Many rodent malaria studies record an aspect of parasite replication (most commonly parasitaemia, which represents the proportion of infected RBC) following an experimental perturbation of the innate immune system. For example, approaches to study the impact of NOS2 (an enzyme involved in producing a reactive free radical) include infecting genetically attenuated mice (13) or mice treated with a small inhibitor drug such as aminoguanidine (14). Such experiments have been conducted on a variety of host genetic backgrounds and using evolutionarily divergent parasite species (i.e. *Plasmodium chabaudi, P. yoelii and P. berghei*) that offer diverse phenotypes. For example, *P. chabaudi* is a model of chronic recrudescing infection whereas *P. berghei* can cause experimental cerebral malaria leading to rapid host mortality (15). Furthermore, these studies also vary in experimental design such as the route and dose of infection. We take advantage of the diversity of these rodent models to collate, curate and statistically analyse infection dynamics when aspects of innate immunity are manipulated, and take a meta-analytic approach to identify and quantify consensus and sources of disagreement among published studies. We then apply meta-regression models to estimate whether host factors (e.g. age, sex), methodology (including adoptive transfer and surgery) or the statistical power of individual studies can shape the impact of immune perturbations on parasite burden (16).

Meta-analyses quantitatively synthesise results from multiple studies using the standard effect size to reveal general patterns in the literature (17). By pooling independent results from multiple studies, meta-analyses boost statistical power (18) and can identify sources of variability across studies to provide biological and epistemological insight (19). Moreover, meta-analyses present a more reproducible and transparent alternative to traditional narrative reviews because conclusions emerge from statistical synthesis of a systematically curated literature (20)Our meta-analysis reveals a small average effect size of experimental manipulation of the innate immune system, which equates to a 6 to 12% change in parasitaemia between control and treatment groups at the peak of infection. Meta-regression models reveal that variation in infection dynamics among studies that use *P. chabaudi* or *P. yoelii* are likely explained by stochasticity (due to small sample sizes) rather than by the effects of experimental perturbations. However, in the *P. berghei* model experimental perturbations have a bigger impact in younger or female mice and an 18% difference in peak parasitaemia can be observed between control and treatment groups when effector molecules (rather than upstream signalling networks) are disrupted.

## 3 Methods

### 3.1 Literature search, eligibility screening and data extraction

Our dataset consists of published articles reporting experimental manipulations of the innate immune system in mice infected with rodent malaria parasites. We identified relevant articles with systematic searches of keywords as well as forward and backward citation searches. For our keyword search, we used the following keywords – innate immunity AND (*Plasmodium* OR rodent malaria) NOT mosquito* NOT human – in four databases (Pubmed, Scopus, Jstor, WoS) in January 2020. We filtered for research articles only and applied no restriction in the year of publication. We screened each article for inclusion in the dataset by first removing duplicates among searches, and then assessed the title and abstract of each article to check the experimental design and results consider a relevant topic. We then assessed the full content (where the full article was available) and retained articles (Supplementary Data 1) that met the set of criteria based on PICO defined elements (21) (stated in Table Supplementary 1) to include their data in the dataset for analysis. We also selected the ten oldest and newest articles that met the criteria for inclusion from the keyword search and conducted forward and backward citation searches to identify and screen relevant articles that are cited by and citing each focal article. We followed the PRISMA ECOEVO 2021 guidelines for reporting systematic reviews and meta-analysis (Figure 1) (22). Overall, from an initial pool of 1507 articles (668 keyword, 460 forward, and 403 backward searches), 84 articles - 47, 20 and 17 from the keyword, backward, and forwards searches, respectively – met the criteria to be included in the dataset for analysis.

**Figure 1.**
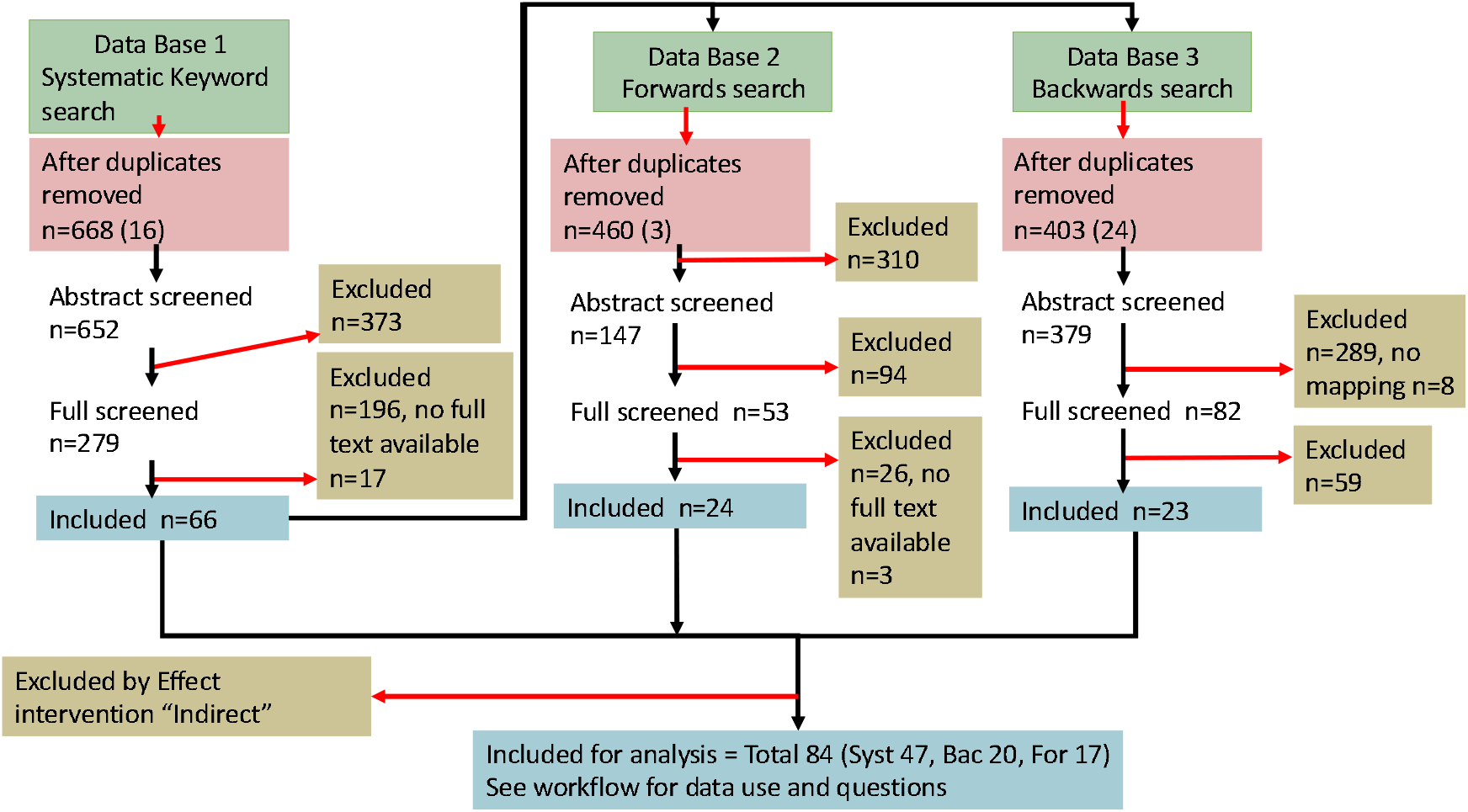
PRISMA flowchart of the literature search and data extraction. Our complete dataset included 84 articles for analysis of innate immune perturbations. Box colours indicate numbers of articles for the literature search (green), screened (pink and purple), included (blue) and excluded studies (yellow), with the number of articles that were removed (red arrows). The following abbreviations refer to: n = number of articles, Syst = Keyword search, Bac = Backwards, For = Forwards.

### 3.2 Data extraction

Our study focuses on analysing patterns of within-host parasite replication under different immune perturbations in rodent malaria infections. The best metric for parasite burden is density (i.e. the number of parasites per μl of blood) but this is rarely reported so instead we extracted parasitaemia (the proportion, or percentage, of infected RBC), and we required studies to report multiple temporally spaced samples for both control and treatment groups. We focus on the acute phase of infection to minimise the influence of other host factors (e.g. RBC availability) that can confound the impacts of immune responses on patterns of parasitaemia (which is affected by the change in the number of RBC). Thus, we include data up to the peak of infection for *P. chabaudi* and non-lethal *P. yoelii*, which we define as the day of infection that either the control or treatment group reaches its maximum reported parasitaemia value. We include all data for *P. berghei* and lethal *P. yoelii* infections because these infections are usually terminated before reaching their peak. We analysed the four infection groups (i.e. *P. chabaudi*, non-lethal and lethal *P. yoelii*, and *P. berghei*) separately as the parasite multiplication rate (and hence infection dynamics) differs markedly between them. When articles included multiple experiments, each suitable experiment was included individually. All data points from figures were extracted using Datathief III v1.7 (2015). AHM conducted all data extraction and TK validated data points. Classification of the immune covariates was determined by AHM and PS.

### 3.3 Effect size – Cohen’s h

We used the Cohen’s h, a standardized effect size, to quantify the difference between proportions (23). We calculated Cohen’s h for each study as the arcsine-transformation of the difference in parasitaemia in the control group compared to that in the treatment group (24). Our study includes experimental manipulations that are expected to augment (e.g. injecting recombinant IFNγ (25)) or attenuate (e.g. depleting phagocytes (26)) innate immunity, which could retard or facilitate parasite replication, respectively. We therefore ignore directionality (27) and use the absolute effect size in all analyses. Put another way, our estimates of Cohen’s h measure the difference between control and treatment groups but do not indicate which group has a higher parasitaemia. For ease of interpretation, we back-transform the meta-analytic mean (conditional upon the control group) to provide an estimate of the overall effect size in terms of the change in parasitaemia at the peak of infection.

### 3.4 Analysis

#### Meta-analysis

We used mixed-effects meta-analytical models and all analyses were conducted in R 4.0.2 (28). We used the rma.mv function, from the metafor package (29), applying the cluster robust var-cov sandwich-type estimator to adjust for small samples and dependent structure. To account for repeated measurements due to multiple effect sizes collected from the same subjects (infections) over time, we included experiment identity as a random effect. We tested for publication bias (whether the published effect sizes were systematically skewed with respect to the overall mean) in our meta-analysis using funnel plots, regression tests and rank tests of the raw effect size. We report heterogeneity among studies as the I^2^ statistic (i.e. as the percentage of variance between effect sizes that cannot be attributed to sampling error due to small sample sizes) (30).

#### Meta-regression

Only the *P. berghei* dataset revealed residual heterogeneity (i.e. variation associated with immune perturbations that cannot be explained by small sample sizes), which allowed us to conduct meta-regression analyses to explore sources of this variation. Moderator information was only available for subsets of data so we carried out a series of univariate meta-regression analyses to examine the moderating effects of host, immunological and experimental design factors on the impacts of immune interventions. Univariate models proved the most appropriate approach due to missing data and/or small sample sizes (Table Supplementary 2) whilst noting that some meta-regressions involve unbalanced samples which may confound the results (16). Meta-regression analysis was performed when the dataset contained more than ten data points (see Table Supplementary 2 for detailed sample sizes across moderator levels). We were unable to assess the effect of host strain because the literature contains too many ambiguous categories within strains to derive a meaningful generalisation. The studied moderators are as follows:

- Host factors
  - Age (centred median age reported in weeks; 5.5 to 11 weeks)
  - Sex (female, male or mixed/unreported)
- Immunological factors (i.e. the target of experimental manipulation)
  - Cell lineage (lymphoid or myeloid cells (or both))
  - Cytokines/chemokines or their receptors
  - Effector function (inflammatory, regulatory or involved in control of cell trafficking)
  - Position in signalling network (receptor to transcription factor (termed input) or downstream targets of transcription factors (termed output))
- Experimental design factors
  - Sampling regime (day post infection)
  - Route of infection (intraperitoneal or intravenous injection of parasites)
  - Infecting dose (the number of parasitised RBC inoculated)
  - Method of immune manipulation (genetic modification (GM), drug administration, surgery, cell transfer or mixed (any combination of these four categories))

We conducted model comparisons using the Akaike Information Criteria corrected for small sample size (AICc) and log-likelihood ratio tests (LRT), and tested for differences between levels of a given moderator using linear hypothesis tests (29).

## 4 Results

### 4.1 Overview of study design

Our meta-analysis began with a systematic literature search. We then assessed the suitability of individual articles for inclusion before data were extracted from suitable studies and effect sizes calculated (Figure 1). The first phase of analysis was to calculate the overall effect size (i.e. meta-analytic mean) for each parasite species using a random-effects meta-analysis. We then estimated heterogeneity across studies to identify potential sources of variation that cannot be attributed to differences in sample size (and which may instead be explained by moderators relating to host factors or experimental design). The second phase of analysis was applied to *P. berghei*, which was the only species for which we detected non-zero heterogeneity. Here we explored the effect of biological and methodological moderators using a series of univariate meta-regressions. Finally, we assessed publication bias graphically and using statistical tests.

### 4.2 Overall effect size

Our meta-analysis found consistent effects of innate immune perturbations on parasite growth dynamics across all three *Plasmodium* species investigated (Figure 2). The overall effect sizes (i.e. the absolute meta-analytic mean Cohen’s h) were 0.1498, 0.2116, 0.2315 and 0.2321 for *P. chabaudi, P. yoelii* lethal, *P. yoelii* non-lethal and *P. berghei*, respectively. These effect sizes are modest as a Cohen’s h of 0.2 and 0.5 are considered a small and medium effect, respectively (23). Back-transforming these effect sizes translates to a change in peak parasitaemia between treatment and control groups of roughly 6 to 12% (6.20%, 8.67%, 9.45% and 11.57% for *P. chabaudi, P. yoelii* lethal, *P. yoelii* non-lethal and *P. berghei*, respectively, Figure 2). Like Cohen’s h, these back transformations are agnostic of the direction of the effects (i.e. they do not report whether parasitaemia was increased or decreased in treatment groups).

**Figure 2.**
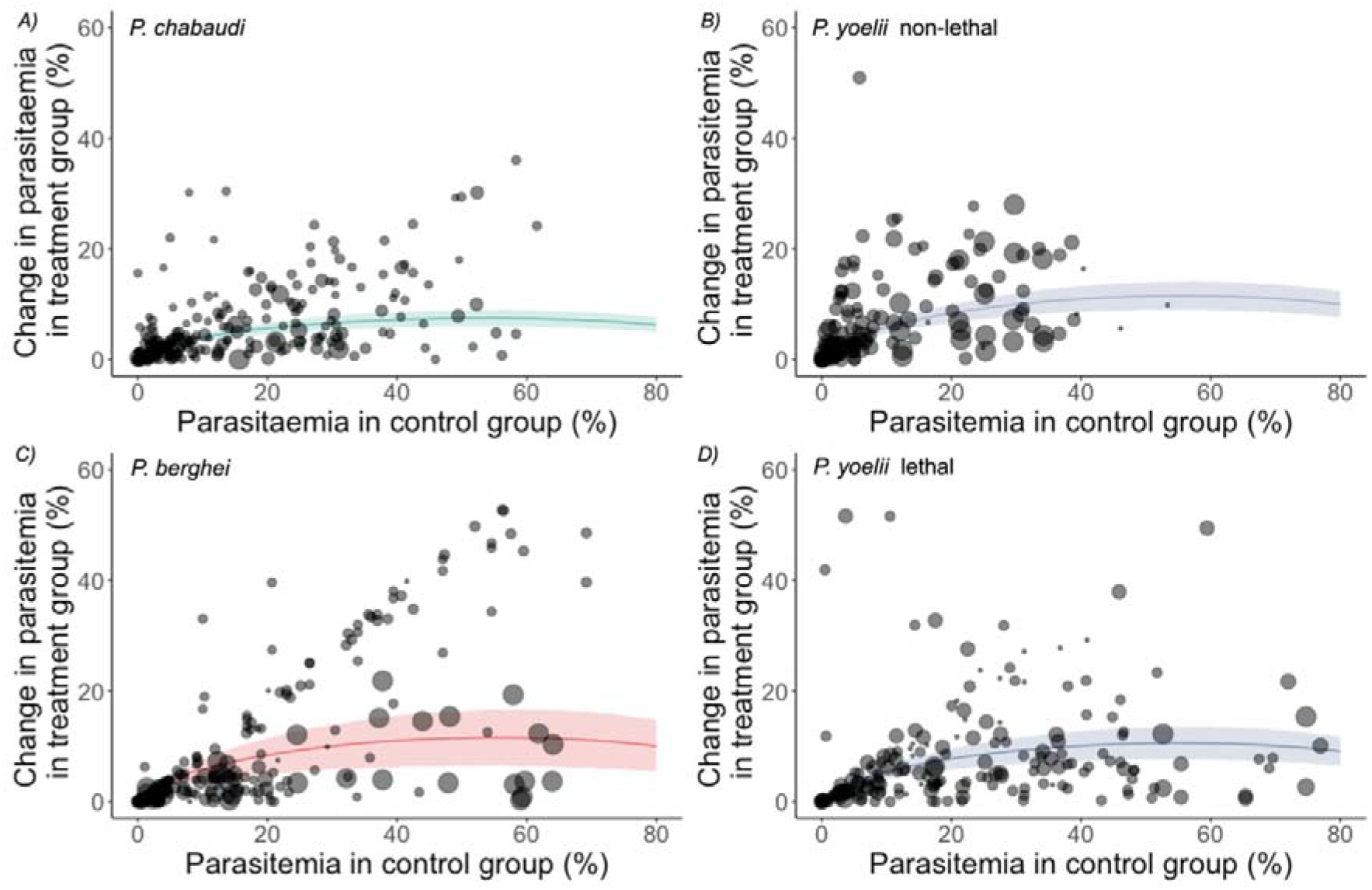
The predicted mean impact of innate immune perturbations from the meta-analytic model. The impact is expressed as the absolute difference in peak parasitaemia in a treatment relative to control group. Note that these plots do not indicate directionality, and so do not indicate that treatment groups have a higher parasitaemia than controls (or vice versa). Instead, they simply illustrate the size of the difference between treatment and control groups. Each panel refers to a different model system: (A) *P. chabaudi*, (B) *P. yoelii* non-lethal, (C) *P. berghei* and (D) *P. yoelii* lethal. Each data point represents experimental sampling points (the size of each dot indicates sample size) and the regression lines and 95% confidence intervals are shown.

### 4.3 Heterogeneity

We detected very low heterogeneity between the effect size of each study for *P. chabaudi, P. yoelii* lethal and *P. yoelii* non-lethal (I^2^ _[total]_= 1.27%, 1.82% and 0.92%, respectively), which is consistent with the notion that random variation due to small sample sizes is responsible for the observed variability (rather than host factors or experimental approaches). In contrast, the *P. berghei* dataset contained comparatively higher heterogeneity (I^2^ _[total]_ = 10.17%), suggesting that factors beyond stochasticity may underpin this variation. We therefore probed for potential sources of variation using a series of univariate meta-regressions, which we report in the following sections (Table 1).

**Table 1.** Comparisons of univariate meta-regression models against the null model, testing the effect of moderators in *P. berghei* infections. Delta AICc, LRT and N refer to differences in Akaike information criteria with correction, likelihood ratio test and the number of studies, respectively.

| Category | Moderator | Delta AICc | LRT | p-value | N |
| --- | --- | --- | --- | --- | --- |
| a) Host factors | Age | 5.5859 | 7.6944 | 0.0055 | 20 |
|  | Sex | 7.5026 | 11.6997 | 0.0029 | 32 |
| b) Immunological factors | Cell lineage | 2.4257 | 1.7713 | 0.4124 | 31 |
|  | Cytokines/chemokines or their receptors | 3.8071 | 0.0000 | 1.0000 | 11 |
|  | Effector function | 3.4694 | 0.7343 | 0.6927 | 29 |
|  | Position in signalling network | 13.3607 | 15.4642 | <.0001 | 24 |
| c) Experimental design factors | Sampling regime | 1.4336 | 3.5208 | 0.0606 | 31 |
|  | Route of infection | 2.0703 | 0.0252 | 0.8738 | 28 |
|  | Infecting dose | 0.4778 | 1.6094 | 0.2046 | 31 |
|  | Method of immune manipulation | 5.6903 | 9.9000 | 0.0071 | 29 |

#### Effect of host traits in *P. berghei* infections

We found that the impact of innate immune perturbations declines with host age (estimate = - 0.1000, SE = 0.0343, t-value = -2.9163, p-value = 0.0092, Figure 3; Table 1). For young mice (at 5.5 weeks) the estimated absolute Cohen’s h was 0.473, which tends toward a medium effect. This constitutes up to a maximum change in peak parasitaemia of ∼18% between treatment and control groups. Furthermore, studies that used only female mice exhibited a stronger impact of immune interventions than studies involving both sexes (or studies that did not report sex) (Figure 3; Table 1). The average absolute Cohen’s h among female mice was 0.396, which is considered a small to medium effect (23) and translates to a maximum difference of ∼16% in peak parasitaemia. However, the lack of experiments using only male mice (N=1) precludes a direct comparison between sexes.

**Figure 3.**
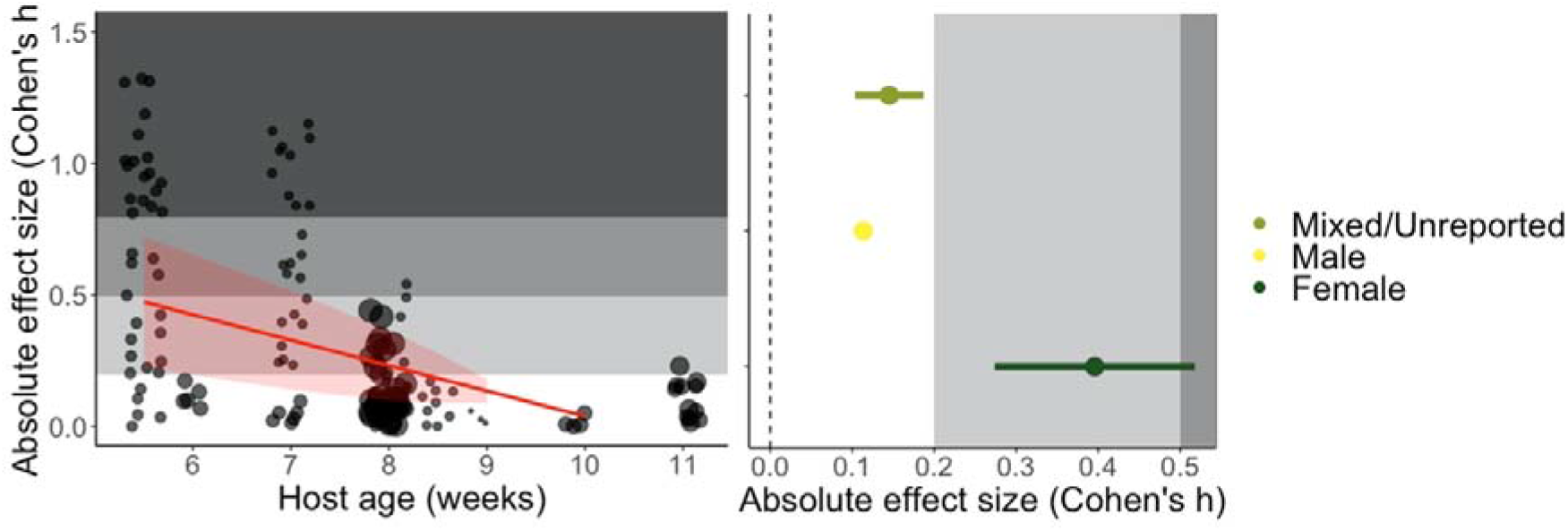
Predicted impact of host (A) age in weeks and (B) sex on the absolute Cohen’s h effect size. a) The regression line is shown in red with a 95% confidence band, and the points represent studies, with point size reflecting the sample size. b) Mean absolute effect ± SEs in three sex categories (females N = 7, males N = 1, unknown N = 23). The grey shading denotes the reference for Cohen’s effect size bands; small (0.2), and medium (0.5) (23).

#### Effect of targeting different aspects of innate immunity in *P. berghei* infections

Out of the four moderators examined only the position of the target within a signalling network showed a significant effect, with interventions that interfere with output (i.e. events downstream of transcription factor signalling) having a significantly larger impact than those modifying input signals (receptor to transcription factor) (F(1, 22) = 16.8373, p-value < 0.001, Figure 4). The average absolute Cohen’s h for output was 0.466, which tends towards a medium effect (23) and translates to a ∼18% difference in peak parasitaemia between treatment and control groups. Other moderators did not significantly explain heterogeneity in the meta-analytic mean and these included: the cell lineage being manipulated (lymphoid versus myeloid); whether interventions targeted cytokines or their receptors; and whether effector molecules had inflammatory versus regulatory functions (Table 1).

**Figure 4.**
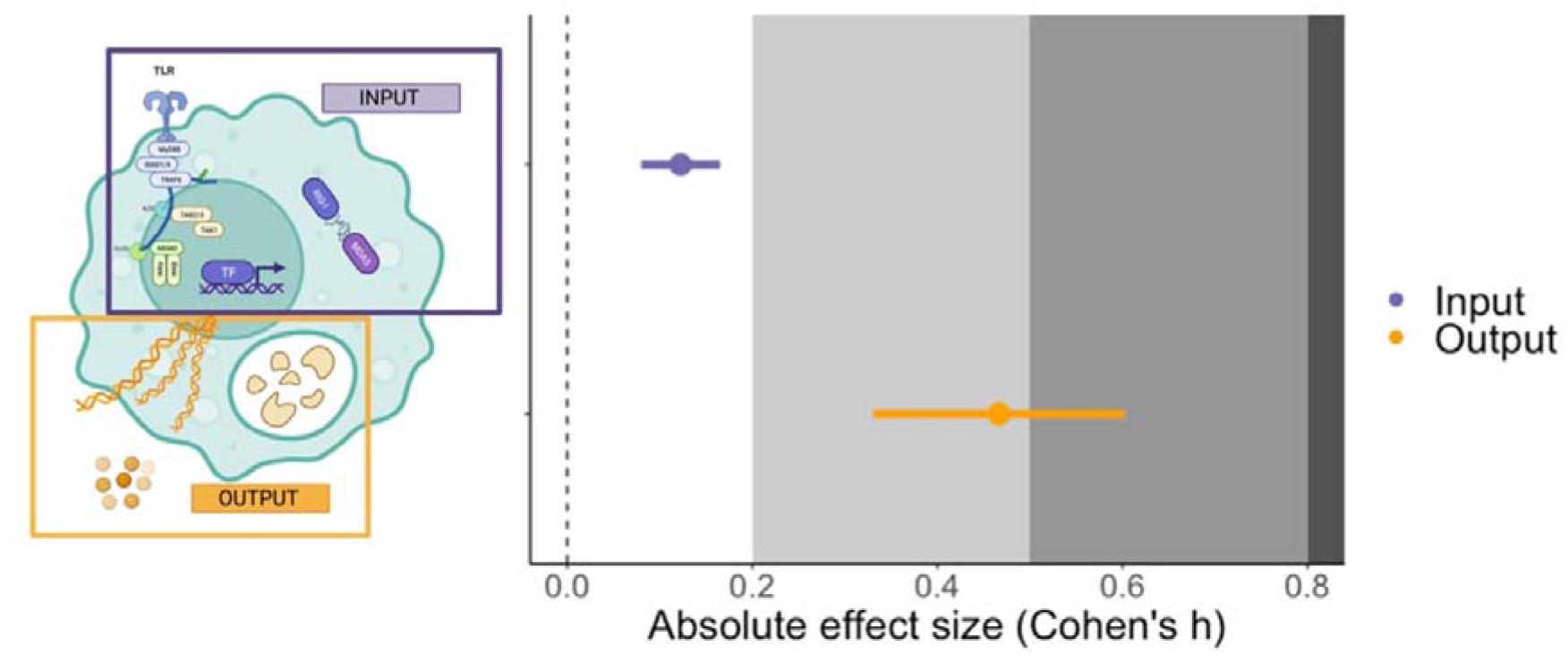
The cartoon (left) illustrates the relative position in an immune signalling network of a target of experimental perturbation: input (purple) refers to factors with an upstream signalling position (e.g. pattern recognition receptors (PRR) and transcription factors) whereas output (orange) refers to downstream effectors (e.g. cytokines, nitric oxide and proteases). The figure (right) shows the mean and ±SE effect size for input and output interventions in the *P. berghei* model. The grey shading denotes the reference for Cohen’s effect size bands; small (0.2), medium (0.5) and large (0.8) (23). The left cartoon was created using BioRender.com.

#### Effect of methodology in *P. berghei* infections

Of the four factors describing different aspects of experimental approaches only the method of immune manipulation explained significant heterogeneity across studies. Drug induced perturbations of the innate immune system (including the administration of recombinant cytokines or monoclonal antibodies as well as small inhibitor molecules) generated the largest impact (Figure 5); the average Cohen’s h was 0.386, which is considered a small to medium effect (23) and translates to a ∼15% difference in parasitaemia between treatment and control groups at the peak of infection. Furthermore, the impact of drug induced perturbations was significantly higher than genetic modification (GM) (χ^2^ = 10.4520, p-value = 0.0012) or experiments that combined more than one methodology (mixed, χ^2^ = 4.561, p-value = 0.0327). Note that it wasn’t possible to estimate the relative effect of adoptive cell transfer or surgery alone because the number of studies that used these techniques was too small. Finally, we found no moderating effects of the sampling regime, route of infection or infecting dose (Table 1).

**Figure 5.**
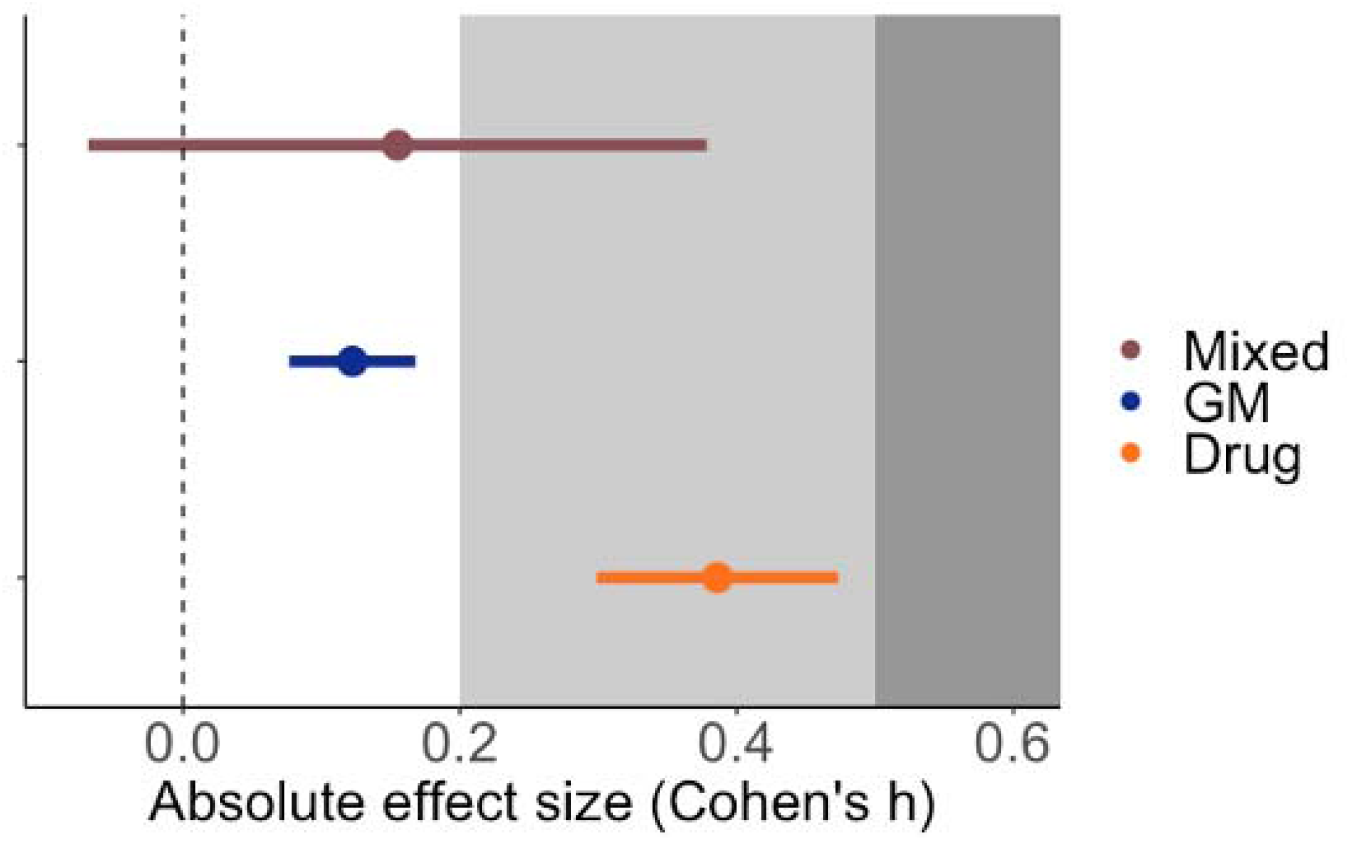
Mean ±SE effect size for the method of immune perturbation in the *P. berghei* model. GM = genetic modification (N = 13), Drug = agonist or antagonist chemical intervention (N = 13) and Mixed = any combination of two or more techniques from genetic modification, drug administration, surgery and adoptive cell transfer (N = 3). The grey shading denotes the reference for Cohen’s effect size bands: small (0.2) and medium (0.5)(23).

### 4.4 Assessment of publication bias

As demonstrated by funnel plots and the regression and rank tests (Figure Supplementary 1 and Table Supplementary 3) we detected little evidence of publication bias in *P. chabaudi, P. yoelii* lethal and *P. yoelii* non-lethal. As we found a potential indication for publication bias in the *P. berghei* dataset (Table Supplementary 3) we carried out a sensitivity analysis by removing the three articles that contributed most to asymmetry. Removing these studies made little difference to the overall mean effect size (Figure Supplementary 2) indicating that our findings are unlikely to be affected by publication bias.

## 5 Discussion

We present a broad ranging meta-analysis of studies spanning four decades of experimental perturbations to identify the role of innate immunity in controlling parasite replication early in infection. Because some studies manipulate immune function in ways that might facilitate parasite replication, whereas others would be expected to decrease parasitaemia, we quantify effect sizes on the absolute scale, which reports the magnitude (not the direction) of the difference between control and treatment groups. Overall, we reveal effect sizes (Cohen’s h) that are considered small for all four datasets, which span three diverse parasite species (Figure 2) (23). The effect sizes range from 0.1498 to 0.2321, which translates to a change in parasitaemia of roughly 6-12% between treatment and control groups at the peak of infection, with *P. berghei* at the upper end of this range. We also reveal that for *P. berghei* the impacts of immune perturbation are larger when younger or female mice are used, and when drugs are administered to manipulate the innate immune system. Furthermore, we observe the greatest effect when effector molecules (or signalling molecules downstream of transcription factors) are targeted (translating to an ∼18% difference in peak parasitaemia). In contrast, there is extremely low heterogeneity across the *P. chabaudi* and *P. yoelii* datasets, which means that variation between studies is better explained by stochasticity (due to small sample sizes) than by host factors or experimental approaches.

The small effect sizes across all datasets have several non-mutually exclusive explanations. These may reflect redundancy in a complex biological system; the innate immune response to malaria is underpinned by a complex network of signalling pathways with considerable functional resilience (31,32). Thus, the innate immune system may compensate and partially restore function following perturbation (33). The observation that effect sizes are larger when effector molecules (the output of signalling networks) are disrupted is consistent with the expectation of greater functional redundancy in upstream events, including receptor binding and signalling. Similarly, the smaller effect sizes observed in older or genetically modified hosts may be due to compensatory mechanisms that can restore immune function through time (34). Thus, the effect sizes we measure may simply reflect the lower bound of the overall effect of innate immunity on parasite replication. However, redundancy and robustness cannot explain the non-significant effects when major aspects of the innate immune system are disrupted or ablated (e.g. following antibody-mediated depletion of the entire myeloid lineage). Thus, an alternative explanation is that innate immunity has only a minor impact on the rate of parasite replication in the first hours and days of infection. That is not to say that innate immune cells do not remove infected red blood cells and free merozoites but rather that the efficiency of this process depends upon help from the adaptive immune system (including CD4^+^ T cells and antibodies), which occurs later in infection (35). In this scenario, the early innate immune response may primarily function to mobilise and recruit phagocytes and lymphocytes to the inflamed spleen to co-ordinate a response that can effectively kill and clear parasites (36). After all, malaria initially triggers interferon-stimulated inflammation and emergency myelopoiesis in the bone marrow (37,38), a conserved innate response that is also induced by viral infection, injury and trauma(39,40); there is no reason to assume that this should be well adapted to early control of malaria parasites. Meta-analysis could establish the importance of mobilisation using complex multivariate analysis that tests for an interaction between time, trafficking and parasite burden but the small number of published studies with relevant data preclude this in the present study (Table Supplementary 2).

We detected low heterogeneity across studies, indicating that the majority of variability is attributable to stochastic variation due to small sample sizes(41). Underpowered studies with small sample sizes are common in *in vivo* animal studies for ethical and logistical considerations (42). However, meta-analyses of studies with small sample sizes are more likely to find low heterogeneity and hence the scope is limited to probe between-study variation (43). As such, larger scale experiments are needed to identify sources of mixed results in the literature, particularly in the *P. chabaudi* and *P. yoelii* models where we detected ∼1% heterogeneity. Why heterogeneity was detected for *P. berghei* (∼10% variation was not attributable to sample size) is unclear. *P. berghei* is not used more often than other parasite species (Table Supplementary 2) and *P. berghei* experiments do not have larger sample sizes, suggesting a biological explanation. It is possible that because *P. berghei* infections are generally short (i.e. terminated before the development of experimental cerebral malaria) there are less opportunities for other factors, such as RBC depletion, to confound the impact of innate immunity on parasite replication. Furthermore, hosts might be more likely to resist than tolerate *P. berghei* compared to other parasite species (12). While *P. berghei* could reveal a greater role for innate immunity in controlling parasite replication it may not reflect malaria more broadly or provide a good model for human infections.

We did not detect a moderating effect of either the timing of sampling or the infectious dose on the impact of immune perturbations. At first glance, an impact of timing may be expected because the possible difference in parasitaemia between control and treatment groups is smaller at the lower boundary. For example, when parasitaemia is low early in infection there is less scope for this to be reduced by immune augmentation as compared to at higher parasitaemias, which are reached later in infection. Fortunately, standard effect sizes, like Cohen’s h, allow for examination of effects free from this constraint, making our inference about the timing of sampling robust (23). The lack of a dose effect was initially unexpected because several mathematical models have previously reported links between the initial parasite density (dose) and immune response to malaria (44–46). Nevertheless, infecting dose does not influence disease severity(47) and perhaps by assuming a strong role for the innate immune response in parasite control we have overlooked the importance of disease tolerance mechanisms (38).

We did, however, identify host sex and age as moderators of innate immunity in the *P. berghei* model; immune perturbations had a more profound impact in young and female mice (Figure 3). This relationship between host sex and innate immunity is consistent with findings that male mice are comparatively immunocompromised (and therefore more susceptible to infection) due to higher testosterone production (48). Yet the role of age in influencing mechanisms of host resistance remains an open question. Immune senescence (leading to reduced functionality) and chronic inflammation (so-called inflammaging) are both hallmarks of the ageing immune system, which could accelerate or slow parasite replication, respectively (49). Nonetheless, it is important to note that our study only includes sexually mature mice (between 5.5 to 11 weeks) and more experimental data are needed to directly examine the impact of innate immunity in older (as well as juvenile and neonatal) mice. As has been remarked elsewhere (50), we noticed that the reporting of host traits tends to be overlooked. For example, many studies in our dataset did not explicitly report the sex (n = 79 out of 140 studies).

In addition to better reporting of host traits, we also advocate for improvements in data reporting. Parasitaemia (the percentage of infected RBC) is widely used as a proxy for parasite burden. However, this metric is problematic for several reasons. Most importantly, the proportion of infected RBC is confounded by the number of circulating RBC, which decreases as infection progresses. Parasites exploit and deplete RBC, and innate immune mechanisms clear RBC (including uninfected cells) and suppress RBC production(51). Thus, longitudinal parasitaemia data provide incomplete information on parasite burden and rates of replication, and it is impossible to distinguish whether top-down (e.g. parasite clearance) or bottom-up (e.g. red blood cell availability) mechanisms underly the observed patterns. Addressing this can be achieved by including RBC counts in addition to parasitaemia at each sampling point. While this is not routine, there is precedent to provide a fuller picture of infection dynamics; among the *P. chabaudi* studies in our data set all but 4 articles reported paired parasitaemia and RBC counts.

### 5.1 Conclusions

Our meta-analysis summarised four decades of research to probe the role of innate immunity in rodent models of malaria. We detected a small overall effect of perturbing the innate immune system on infection dynamics and revealed that manipulating effector molecules in young female mice by administering drugs (including monoclonal antibodies) has the largest impact on the replication of *P. berghei*. However, small sample sizes likely explain much of the variation across studies in the *P. chabaudi* and *P. yoelii* models. Why the impact of different experimental approaches varies between parasite species is unclear but raises questions about how to translate findings from rodent models to human malaria. Important differences exist between parasite species that infect humans and rodents, including that rodent *Plasmodium spp*. complete asexual replication in half the time it takes for the major human parasites (*P. falciparum* and *P. vivax)*. Perhaps the capacity for rodent parasites to replicate to much higher densities causes resource limitation to play a larger role in limiting parasite burden. No study system marries ecological realism, translational value, and tractability. Yet, perhaps the rapid development of human challenge models allows for the role of innate immunity early in infection to be tested experimentally by integrating human data with mathematical modelling to test long-standing (and alternative) hypotheses. Certainly, by combining a more integrative approach with larger scale experiments that report more comprehensive data should help resolve conflicting reports on the role of innate immunity in malaria.

## Acknowledgements

We thank all the authors contributing through primary data.

## Availability of data and materials

The datasets are available in the Edinburgh DataShare repository: https://datashare.ed.ac.uk/handle/10283/3204 URL- reecelab

## Authors’ contributions

Conceptualization, all authors; Analysis, AHM and TK; Writing-Original Draft, AHM, TK, SER; Writing-Review and Editing, all authors; Supervision, TK, SER.

## References

1. Graham AL, Tate AT. Are we immune by chance?. Elife. (2017) 27:6.

2. Götz A, Ty M, Chora AF, Zuzarte-Luís V, Mota MM, Rodriguez A. Innate Immunity to Malaria. In: Mota, M., Rodriguez, A. (eds) Malaria. Springer, Cham. (2017) 1;3–25. https://doi.org/10.1007/978-3-319-45210-4_1

3. Stevenson MM, Riley EM. Innate immunity to malaria. Nat Rev Immunol. (2004) 4(3):169–80.

4. Perez-Mazliah D, Langhorne J. CD4 T-cell subsets in malaria: Th1/Th2 revisited. Front Immunol. Frontiers Media S.A. (2015) 12;5:671.

5. Sebina I, Haque A. Effects of type I interferons in malaria. Immunology (2018) 155;2:176–85.

6. Clark IA, Chaudhri G. Tumour necrosis factor may contribute to the anaemia of malaria by causing dyserythropoiesis and erythrophagocytosis. Br J Haematol (1988) 70(1):99–103.

7. Belyaev NN, Brown DE, Diaz AIG, Rae A, Jarra W, Thompson J, et al. Induction of an IL7-R+c-Kithi myelolymphoid progenitor critically dependent on IFN-γ signaling during acute malaria. Nat Immunol (2010) 11(6):477–85.

8. Kirkman LA, Deitsch KW. Vive la différence: Exploiting the differences between rodent and human malarias. Trends Parasitol. (2020) 36:504–11.

9. Stephens R, Culleton RL, Lamb TJ. The contribution of Plasmodium chabaudi to our understanding of malaria. Trends Parasitol. (2012) 28(2):73–82.

10. Chavatte JM, Snounou G. Controlled human malaria infection—maker and breaker of dogma. PLoS Med. (2021) 18;4:e1003591.

11. Brown AC, Guler JL. From circulation to cultivation: Plasmodium in vivo versus in vitro. Trends Parasitol. (2020) 36;11:914–26.

12. Olatunde AC, Cornwall DH, Roedel M, Lamb TJ. Mouse models for unravelling immunology of blood stage malaria. Vaccines (Basel). (2022) 10;9:1525.

13. Green SJ, Scheller LF, Marletta MA, Seguin MC, Klotz FW, Slayter M, et al. Nitric oxide: cytokine-regulation of nitric oxide in host resistance to intracellular pathogens. Immunol Lett. (1994) 43:87–94.

14. Van der Heyde HC, Gu Y, Zhang Q, Sun G, Grisham MB. Nitric oxide is neither necessary nor sufficient for resolution of Plasmodium chabaudi malaria in mice. J Immunol. (2000) 165;6:3317–23.

15. Brian De Souza J, Hafalla JCR, Riley EM, Couper KN. Cerebral malaria: Why experimental murine models are required to understand the pathogenesis of disease. Parasitology. (2010) 137;5:755–72.

16. Harrer M, Cuijpers P, Furukawa TA, Ebert DD. Doing meta-analysis with R: A hands-on guide. 1st ed. Boca Raton, FL and London: Chapman & Hall/CRC Press. (2021).

17. Gurevitch J, Nakagawa S. Gurevitch Jessica, and Shinichi Nakagawa, ‘Research synthesis methods in ecology’, in Gordon A. Fox, Simoneta Negrete-Yankelevich, and Vinicio J. Sosa (eds), Ecological Statistics: Contemporary theory and application. Oxford, (2015) Oxford Academic.

18. Cohn LD, Decker BJ. How meta-analysis increases statistical power. Psychol Methods. (2003) 8;3:243–53.

19. Baker WL, Michael White C, Cappelleri JC, Kluger J, Coleman CI. Understanding heterogeneity in meta-analysis: the role of meta-regression. Int J Clin Pract. (2009) 63;10:1426–34.

20. Garg AX, Hackam D, Tonelli M. Systematic review and meta-analysis: When one study is just not enough. Clin J Am Soc Nephrol. (2008) 3;1:253–60.

21. McKenzie J, Brennan S, Ryan R, Thomson H, Johnston R, Thomas J. Chapter 3 Defining the criteria for including studies and how they will be grouped for the synthesis. (2022). In: Higgins J, Thomas J, Chandler J, Cumpston M, Li T, Page M, et al., editors. Cochrane handbook for systematic reviews of interventions version 6.3. [Accessed June 2022]. https://www.training.cochrane.org/handbook.

22. O’Dea RE, Lagisz M, Jennions MD, Koricheva J, Noble DWA, Parker TH, et al. Preferred reporting items for systematic reviews and meta-analyses in ecology and evolutionary biology: a <scp>PRISMA</scp> extension. Biol Rev. (2021) 96;5:1695–722.

23. Cohen J. Statistical power analysis for the behavioral sciences. Saint Louis, United States: Elsevier Science & Technology. (1977). p.179–214

24. Rosenthal R. Parametric measures of effect size. In: Cooper H, Hedges LV, editors. The handbook of research synthesis. New York: Russell Sage Foundation (1994). p. 231–

25. Clark IA, Hunt NH, Butcher GA, Cowden WB. Inhibition of murine malaria (Plasmodium chabaudi) in vivo by recombinant interferon-γ or tumor necrosis factor, and its enhancement by butylated hydroxyanisole. J. Immunol. (1987) 139;10:3493–6.

26. Terkawi MA, Nishimura M, Furuoka H, Nishikawa Y. Depletion of phagocytic cells during nonlethal Plasmodium yoelii infection causes severe malaria characterized by acute renal failure in mice. Infect Immun. (2016) 11;84(3):845–55.

27. Atallah MB, Tandon V, Hiam KJ, Boyce H, Hori M, Atallah W, et al. ImmunoGlobe: Enabling systems immunology with a manually curated intercellular immune interaction network. BMC Bioinformatics. (2020) 21;1:1–18.

28. R Core Team. R: A Language and Environment for Statistical Computing. Vienna, Austria (2020).

29. Viechtbauer W. metafor: Meta-analysis package for R. (2021)

30. Higgins JPT, Thompson SG, Deeks JJ, Altman DG. Measuring inconsistency in meta-analyses. BMJ (2003) 327;7414:557–60.

31. Gazzinelli RT, Kalantari P, Fitzgerald KA, Golenbock DT. Innate sensing of malaria parasites. Nat Rev Immunol. (2014) 14. p. 744–57.

32. Gowda DC, Wu X. Parasite recognition and signaling mechanisms in innate immune responses to malaria. Front Immunol. (2018) 9: 3006.

33. Nish S, Medzhitov R. Host Defense Pathways: role of redundancy and compensation in infectious disease phenotypes. Immunity. (2011) 34;5:629.

34. El-Brolosy MA, Stainier DYR. Genetic compensation: A phenomenon in search of mechanisms. PLoS Genetics. (2017) 13.

35. Good MF, Doolan DL. Immune effector mechanisms in malaria. Curr Opin Immunol. (1999) 11;4:412–9.

36. Engwerda CR, Beattie L, Amante FH. The importance of the spleen in malaria. Trends Parasitol (2005) 21(2):75–80.

37. Spaulding E, Fooksman D, Moore JM, Saidi A, Feintuch CM, Reizis B, et al. STING-licensed macrophages prime type I IFN production by plasmacytoid dendritic cells in the bone marrow during severe Plasmodium yoelii malaria. PLoS Pathog (2016) 12(10)

38. Nahrendorf W, Ivens A, Spence PJ. Inducible mechanisms of disease tolerance provide an alternative strategy of acquired immunity to malaria. Elife. (2021) 10.

39. Kelly LS, Darden DB, Fenner BP, Efron PA, Mohr AM. The Hematopoietic stem/progenitor cell response to hemorrhage, injury, and sepsis: A review of pathophysiology. Shock (2021) 56(1):30–41.

40. Hawthorne AL, Popovich PG. Emerging concepts in myeloid cell biology after spinal cord injury. Neurotherapeutics (2011) 8(2):252–61.

41. Higgins JPT, Thompson SG, Deeks JJ, Altman DG. Measuring inconsistency in meta-analyses. BMJ (2003) 327(7414):557–60.

42. Vesterinen HM, Sena ES, Egan KJ, Hirst TC, Churolov L, Currie GL, et al. Meta-analysis of data from animal studies: A practical guide. J. Neurosci. Methods. (2014) 221:92–102. 39.

43. Rücker G, Schwarzer G, Schumacher M, Carpenter J. Are large trials less reliable than small trials?. J Clin Epidemiol. (2009) 62;8:886–7.

44. Kamiya, T, Greischar, MA, Schneider, DS, Mideo, N. Uncovering drivers of dose-dependence and individual variation in malaria infection outcomes. PLoS Comput. Biol. (2020) 16(10), e1008211. https://doi.org/10.1371/JOURNAL.PCBI.1008211

45. Metcalf, CJE, Graham, AL, Huijben, S, Barclay, VC, Long, GH, Grenfell, et al. Partitioning and quantifying regulatory mechanisms acting on within-host malaria using the effective propagation number. Science. (2011) 333(6045):984. https://doi.org/10.1126/SCIENCE.1204588

46. Camponovo F, Lee TE, Russell JR, Burgert L, Gerardin J, Penny MA. Mechanistic within-host models of the asexual Plasmodium falciparum infection: a review and analytical assessment. Malar J. (2021) 20;1:1–22.

47. Glynn JR, Collins WE, Jeffery GM, Bradley DJ. Infecting dose and severity of falciparum malaria. Trans R Soc Trop Med Hyg (1995) 89(3):281–3.

48. Klein SL. Hormonal and immunological mechanisms mediating sex differences in parasite infection. Parasite Immunol. (2004) 26;6–7:247–64.

49. Borgoni S, Kudryashova KS, Burka K, de Magalhães JP. Targeting immune dysfunction in aging. Ageing Res Rev (2021) 70.

50. Vom Steeg LG, Flores-Garcia Y, Zavala F, Klein SL. Irradiated sporozoite vaccination induces sex-specific immune responses and protection against malaria in mice. Vaccine. (2019) 37;32:4468–76.

51. White NJ. Anaemia and malaria. Malar J (2018) 17(1).

