## supplementary for "The impact of innate immunity on malaria parasite infection dynamics"

### 1 Supplementary Figures and Tables

#### 1.1 Supplementary Figures

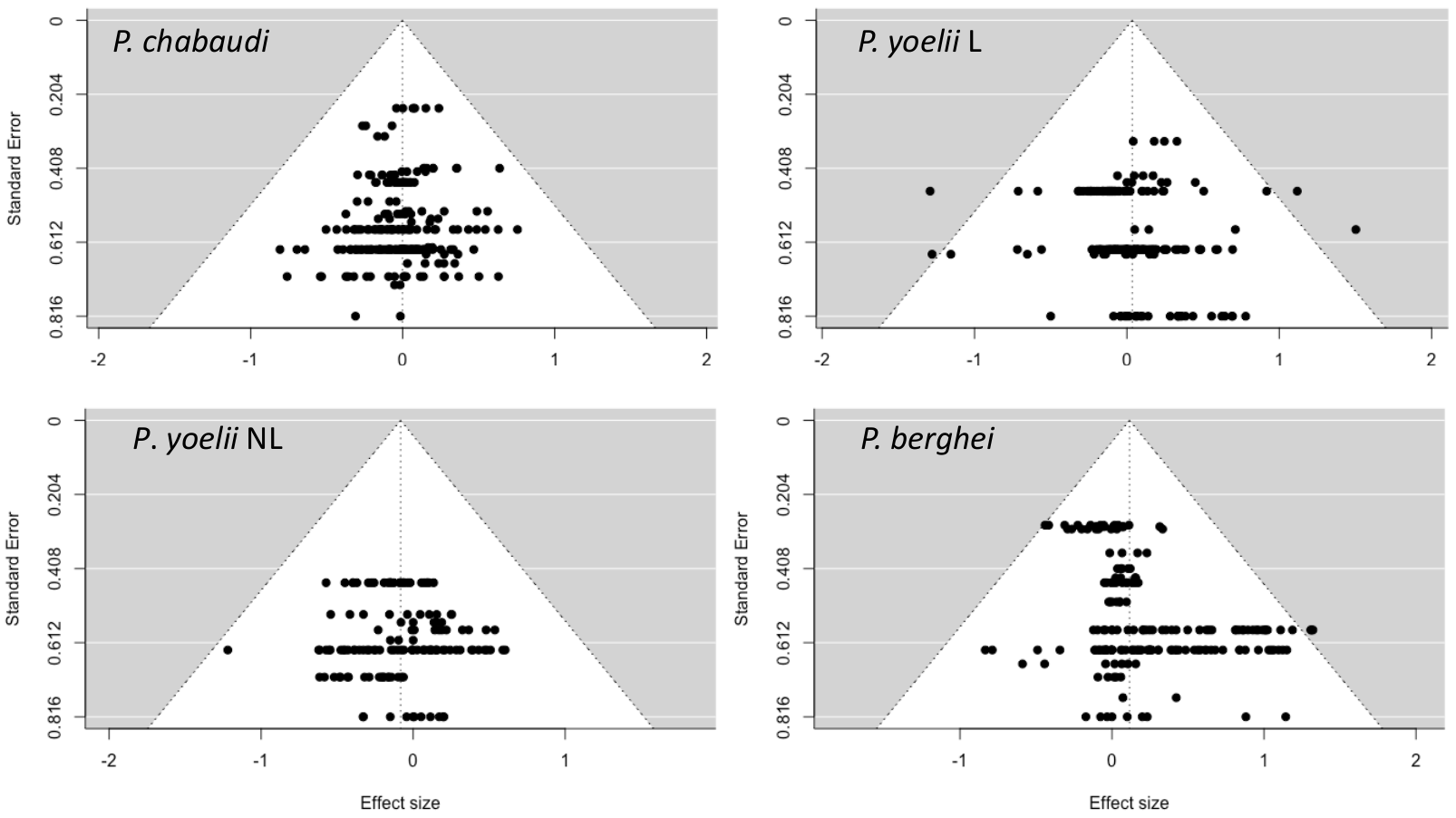

Supplementary Figure 1. Funnel plots showing the standard error (y-axis) against raw effect size (Cohen’s; x-axis). The distribution of studies with high precision (near the top) is expected close to the average in the absence of publication bias, while those with low precision (near the bottom) are expected to be distributed more widely on both sides. The grey area indicates a significant departure from this expectation with a 95% confidence level.

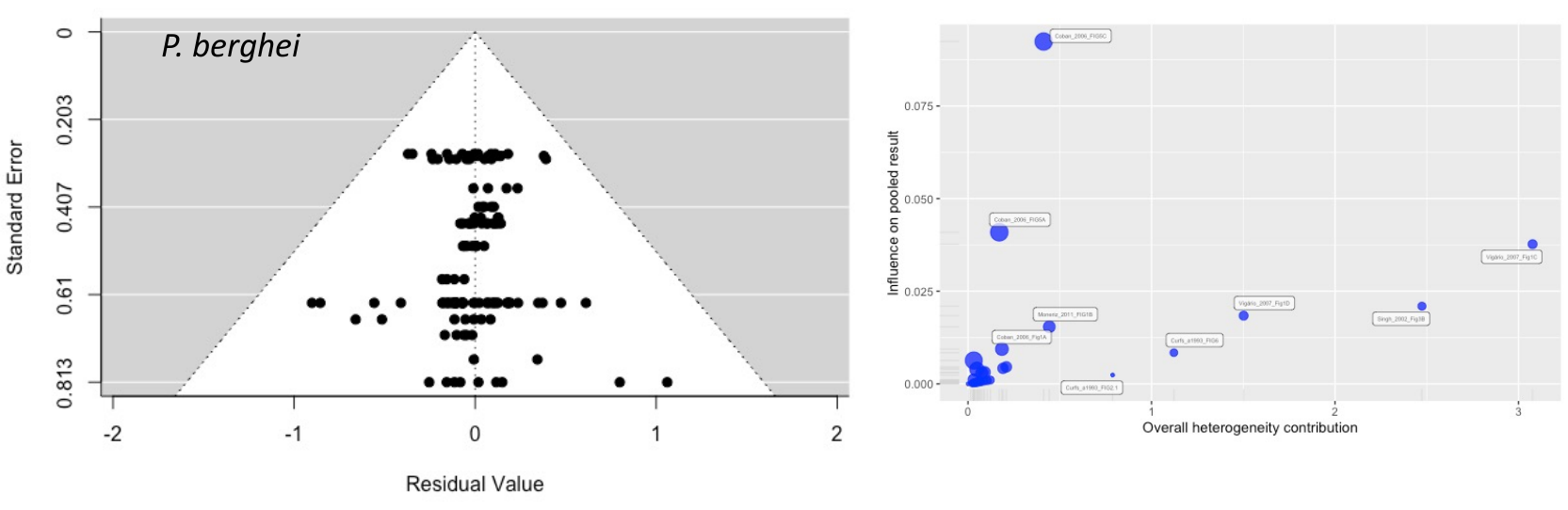

Supplementary Figure 2. Modified funnel plot of *P. berghei* data after removing three influential articles. “Boujat” graph showing mean effect size per study (right), the three articles removed correspond to the articles with higher influence and heterogeneity contribution. After omitting these articles regression to inverse variance is not significant, Egger’s test, intercept = 0.1038, se=0.0854, t=1.2151, df=127, p=0.2266, 95%CI -0.0652- 0.2728.

#### 1.2 Supplementary Tables

**Table supplementary 1.** Explicit exclusion criteria. Table displays the eligibility criteria employed based on PICO defined elements (McKenzie et al., 2022).

|  | Explicit exclusion criteria |
| --- | --- |
| Subject | The subject is not related to the question of how innate immune components impact on parasite burden.  Indirectly testing the effect: correlative or speculative mechanistic link to immune function.  Experiments have no control and treatment groups (for example host strains labelled as susceptible vs resistant) |
| Type | Not a research article (review, opinion, protocol, conference report).  Article used data from another in the primary literature  Not in English |
| Comparator | Uses parasite species that is not one of *P. berghei, yoelii, chabaudi, vinckei*  Uses a non-mouse host species.  Infections initiated from a phase other than the blood stage (liver or mosquito derived infections)  Co-infected or multiply infected hosts  Uses genetically modified parasites (GMO) with a replication rate phenotype, or a mix of parasite antigens that activate the immune response, or attenuated parasites.  Uses hosts previously challenged with *Plasmodium* parasites or their antigens |
| Intervention | Data collected from *in vitro* or *ex vivo* approaches  Not directly testing a component(s) of the innate immune response (e.g. by perturbing adaptive immune function) |
| Response | No quantification of parasite dynamics during infections (e.g. only point estimate of difference in parasite performance between treatment and control)  Day post infection information not included for parasite performance estimates  Data collected after the perturbation is expected to have exerted its influence on parasite performance  No justification given to relate how the sampling regime captures parasite dynamics of control and treatment groups  Data not suitable to relate to parasitaemia |

Table supplementary 2. The table shows the number of studies per moderator variables. Labels for levels within moderators: position in signalling network (in = input, out = output), cytokines/chemokines or their receptors (cc = cytokines/chemokines, rec = receptors), effector function (inflam = inflammatory, reg = regulatory, traff = cell trafficking), method of immune manipulation (GM = genetic modification) and route of infection (Ip = intraperitoneal injection, Iv = intravenous injection).

|  | Immune factors | | | | Methodology | | Host factors | |
| --- | --- | --- | --- | --- | --- | --- | --- | --- |
| Parasite | Position in signalling network | Cytokines/  chemokines or their receptors | Effector function | Cell lineage | Method of immune manipulation | Route of infection | Sex | Age |
| Pc | 46 | 37 | 47 | 57 | 61 | 43 | 61 | 52 |
|  | 18 in  28 out | 26 cc  11rec | 21 inflam  26 reg | 9 lymphoid  15 myeloid  33 both | 5 mixed  27 drug  29 GM | 37 Ip  6 Iv | 4 male  21 female | NA |
| Pb | 24 | 11 | 29 | 31 | 29 | 28 | 32 | 20 |
|  | 19 in  5 out | 10cc  1 rec | 16 inflam  12 reg  1 traff | 1 lymphoid  7 myeloid  23 both | 3 mixed  13 drug  13 GM | 25 Ip  3 Iv | 1 male  7 female | NA |
| PyL | 19 | 10 | 17 | 23 | 23 | 22 | 23 | 22 |
|  | 12 in  7 out | 7 cc  3 rec | 10 inflam  3 reg  4 traff | 1 lymphoid  2 myeloid  20 both | 1 mixed  8 drug  14 GM | 7 Ip 15 Iv | 1 male  18 female | NA |
| PyNL | 16 | 8 | 17 | 23 | 24 | 22 | 24 | 20 |
|  | 11 in  5 out | 4 cc  4 rec | 10 inflam  7 reg | 1 lymphoid  6 myeloid  16 both | 1 mixed  13 drug  10 GM | 12 Ip 10 Iv | 1 male  8 female | NA |

Table supplementary 3. Table shows funnel plot asymmetry. Two common meta-analytic tests were conducted to assess publication bias (Rank tests and Egger’s test).

|  | Rank correlation test for funnel plot asymmetry | Egger’s test (regression to inverse variance) |
| --- | --- | --- |
| *P. chabaudi* | Kendall's tau=-0.0022, p-value=0.9612 | Intercept=-0.0212, SE=0.0518, t=-0.4100, df247, p-value=0.6821 |
| *P. berghei* | Kendall's tau=0.2051, p-value < .0001 | Intercept=0.2747, SE=0.1011, t=2.7180, df188, p-value=0.0072 |
| *P. yoelii* lethal | Kendall's tau=0.2051, p-value < .0001 | Intercept=0.0454, SE=0.1276, t=0.3557, df190, p-value=0.7225 |
| *P. yoelii* non-lethal | Kendall's tau=-0.0438, p-value=0.4827 | Intercept=-0.0194, SE=0.1466, t=-0.1320, df137, p-value=0.8952 |

### 2. Supplementary Data

#### 2.1 Supplementary Data 1

List of references of articles contributing with data for the meta-analyses. Associated effect sizes and metadata can be found in Edinburgh DataShare repository.

Abo, T., & Sekikawa, H. (2002). Extrathymic T cells in malaria protection, including evidence for the onset of erythropoiesis in the liver during infection. *Archives of histology and cytology*, *65*(2), 127–132. <https://doi.org/10.1679/aohc.65.127>

Ataide, M. A., Andrade, W. A., Zamboni, D. S., Wang, D., Souza, M.doC., Franklin, B. S., Elian, S., Martins, F. S., Pereira, D., Reed, G., Fitzgerald, K. A., Golenbock, D. T., & Gazzinelli, R. T. (2014). Malaria-induced NLRP12/NLRP3-dependent caspase-1 activation mediates inflammation and hypersensitivity to bacterial superinfection. *PLoS pathogens*, *10*(1), e1003885. <https://doi.org/10.1371/journal.ppat.1003885>

Bakir, H. Y., Tomiyama-Miyaji, C., Watanabe, H., Nagura, T., Kawamura, T., Sekikawa, H., & Abo, T. (2006). Reasons why DBA/2 mice are resistant to malarial infection: expansion of CD3int B220+ gammadelta T cells with double-negative CD4- CD8- phenotype in the liver. *Immunology*, *117*(1), 127–135. <https://doi.org/10.1111/j.1365-2567.2005.02273.x>

Bastos, K. R., Barboza, R., Elias, R. M., Sardinha, L. R., Grisotto, M. G., Marinho, C. R., Amarante-Mendes, G. P., Alvarez, J. M., & Lima, M. R. (2002). Impaired macrophage responses may contribute to exacerbation of blood-stage *Plasmodium chabaudi chabaudi* malaria in interleukin-12-deficient mice. *Journal of interferon & cytokine research: the official journal of the International Society for Interferon and Cytokine Research*, *22*(12), 1191–1199. <https://doi.org/10.1089/10799900260475713>

Batchelder, J. M., Burns, J. M., Jr, Cigel, F. K., Lieberg, H., Manning, D. D., Pepper, B. J., Yañez, D. M., van der Heyde, H., & Weidanz, W. P. (2003). *Plasmodium chabaudi adami*: interferon-gamma but not IL-2 is essential for the expression of cell-mediated immunity against blood-stage parasites in mice. *Experimental parasitology*, *105*(2), 159–166. <https://doi.org/10.1016/j.exppara.2003.12.003>

Borges da Silva, H., Fonseca, R., Cassado, A.dosA., Machado de Salles, É., de Menezes, M. N., Langhorne, J., Perez, K. R., Cuccovia, I. M., Ryffel, B., Barreto, V. M., Marinho, C. R., Boscardin, S. B., Álvarez, J. M., D'Império-Lima, M. R., & Tadokoro, C. E. (2015). In vivo approaches reveal a key role for DCs in CD4+ T cell activation and parasite clearance during the acute phase of experimental blood-stage malaria. *PLoS pathogens*, *11*(2), e1004598. <https://doi.org/10.1371/journal.ppat.1004598>

Campanella, G. S., Tager, A. M., El Khoury, J. K., Thomas, S. Y., Abrazinski, T. A., Manice, L. A., Colvin, R. A., & Luster, A. D. (2008). Chemokine receptor CXCR3 and its ligands CXCL9 and CXCL10 are required for the development of murine cerebral malaria. *Proceedings of the National Academy of Sciences of the United States of America*, *105*(12), 4814–4819. <https://doi.org/10.1073/pnas.0801544105>

Choudhury, H. R., Sheikh, N. A., Bancroft, G. J., Katz, D. R., & De Souza, J. B. (2000). Early nonspecific immune responses and immunity to blood-stage nonlethal *Plasmodium yoelii* malaria. *Infection and immunity*, *68*(11), 6127–6132. <https://doi.org/10.1128/IAI.68.11.6127-6132.2000>

Clark, I. A., & Hunt, N. H. (1983). Evidence for reactive oxygen intermediates causing hemolysis and parasite death in malaria. *Infection and immunity*, *39*(1), 1–6. https://doi.org/10.1128/iai.39.1.1-6.1983

Coban, C., Ishii, K. J., Uematsu, S., Arisue, N., Sato, S., Yamamoto, M., Kawai, T., Takeuchi, O., Hisaeda, H., Horii, T., & Akira, S. (2007). Pathological role of Toll-like receptor signaling in cerebral malaria. *International immunology*, *19*(1), 67–79. <https://doi.org/10.1093/intimm/dxl123>

Couper, K. N., Blount, D. G., Hafalla, J. C., van Rooijen, N., de Souza, J. B., & Riley, E. M. (2007). Macrophage-mediated but gamma interferon-independent innate immune responses control the primary wave of *Plasmodium yoelii* parasitemia. *Infection and immunity*, *75*(12), 5806–5818. <https://doi.org/10.1128/IAI.01005-07>

Curfs, J. H., Hermsen, C. C., Kremsner, P., Neifer, S., Meuwissen, J. H., Van Rooyen, N., & Eling, W. M. (1993). Tumour necrosis factor-alpha and macrophages in *Plasmodium berghei*-induced cerebral malaria. *Parasitology*, *107 ( Pt 2)*, 125–134. <https://doi.org/10.1017/s0031182000067226>

Duan, X., Imai, T., Chou, B., Tu, L., Himeno, K., Suzue, K., Hirai, M., Taniguchi, T., Okada, H., Shimokawa, C., & Hisaeda, H. (2013). Resistance to malaria by enhanced phagocytosis of erythrocytes in LMP7-deficient mice. *PloS one*, *8*(3), e59633. <https://doi.org/10.1371/journal.pone.0059633>

Edwards, C. L., Best, S. E., Gun, S. Y., Claser, C., James, K. R., de Oca, M. M., Sebina, I., Rivera, F.deL., Amante, F. H., Hertzog, P. J., Engwerda, C. R., Renia, L., & Haque, A. (2015). Spatiotemporal requirements for IRF7 in mediating type I IFN-dependent susceptibility to blood-stage Plasmodium infection. *European journal of immunology*, *45*(1), 130–141. <https://doi.org/10.1002/eji.201444824>

Elased, K. M., Taverne, J., & Playfair, J. H. (1996). Malaria, blood glucose, and the role of tumour necrosis factor (TNF) in mice. *Clinical and experimental immunology*, *105*(3), 443–449. <https://doi.org/10.1046/j.1365-2249.1996.d01-781.x>

Elased, K., De Souza, J. B., & Playfair, J. H. (1995). Blood-stage malaria infection in diabetic mice. *Clinical and experimental immunology*, *99*(3), 440–444. <https://doi.org/10.1111/j.1365-2249.1995.tb05570.x>

Feng, Y., Zhu, X., Wang, Q., Jiang, Y., Shang, H., Cui, L., & Cao, Y. (2012). Allicin enhances host pro-inflammatory immune responses and protects against acute murine malaria infection. *Malaria journal*, *11*, 268. <https://doi.org/10.1186/1475-2875-11-268>

Fernandes, E. S., Brito, C. X., Teixeira, S. A., Barboza, R., dos Reis, A. S., Azevedo-Santos, A. P., Muscará, M., Costa, S. K., Marinho, C. R., Brain, S. D., & Grisotto, M. A. (2014). TRPV1 antagonism by capsazepine modulates innate immune response in mice infected with *Plasmodium berghei* ANKA. *Mediators of inflammation*, *2014*, 506450. <https://doi.org/10.1155/2014/506450>

Finney, C. A., Lu, Z., Hawkes, M., Yeh, W. C., Liles, W. C., & Kain, K. C. (2010). Divergent roles of IRAK4-mediated innate immune responses in two experimental models of severe malaria. *The American journal of tropical medicine and hygiene*, *83*(1), 69–74. <https://doi.org/10.4269/ajtmh.2010.09-0753>

Franklin, B. S., Rodrigues, S. O., Antonelli, L. R., Oliveira, R. V., Goncalves, A. M., Sales-Junior, P. A., Valente, E. P., Alvarez-Leite, J. I., Ropert, C., Golenbock, D. T., & Gazzinelli, R. T. (2007). MyD88-dependent activation of dendritic cells and CD4(+) T lymphocytes mediates symptoms, but is not required for the immunological control of parasites during rodent malaria. *Microbes and infection*, *9*(7), 881–890. <https://doi.org/10.1016/j.micinf.2007.03.007>

Furuta, T., Imajo-Ohmi, S., Fukuda, H., Kano, S., Miyake, K., & Watanabe, N. (2008). Mast cell-mediated immune responses through IgE antibody and Toll-like receptor 4 by malarial peroxiredoxin. *European journal of immunology*, *38*(5), 1341–1350. <https://doi.org/10.1002/eji.200738059>

Geurts, N., Martens, E., Verhenne, S., Lays, N., Thijs, G., Magez, S., Cauwe, B., Li, S., Heremans, H., Opdenakker, G., & Van den Steen, P. E. (2011). Insufficiently defined genetic background confounds phenotypes in transgenic studies as exemplified by malaria infection in Tlr9 knockout mice. *PloS one*, *6*(11), e27131. <https://doi.org/10.1371/journal.pone.0027131>

Gowda, N. M., Wu, X., & Gowda, D. C. (2012). TLR9 and MyD88 are crucial for the development of protective immunity to malaria. *Journal of immunology (Baltimore, Md. : 1950)*, *188*(10), 5073–5085. <https://doi.org/10.4049/jimmunol.1102143>

Gramaglia, I., Velez, J., Combes, V., Grau, G. E., Wree, M., & van der Heyde, H. C. (2017). Platelets activate a pathogenic response to blood-stage *Plasmodium* infection but not a protective immune response. *Blood*, *129*(12), 1669–1679. <https://doi.org/10.1182/blood-2016-08-733519>

Grun, J. L., & Weidanz, W. P. (1981). Immunity to *Plasmodium chabaudi adami* in the B-cell-deficient mouse. *Nature*, *290*(5802), 143–145. <https://doi.org/10.1038/290143a0>

Grun, J. L., Long, C. A., & Weidanz, W. P. (1985). Effects of splenectomy on antibody-independent immunity to *Plasmodium chabaudi adami* malaria. *Infection and immunity*, *48*(3), 853–858. <https://doi.org/10.1128/iai.48.3.853-858.1985>

Hafalla, J. C., Burgold, J., Dorhoi, A., Gross, O., Ruland, J., Kaufmann, S. H., & Matuschewski, K. (2012). Experimental cerebral malaria develops independently of caspase recruitment domain-containing protein 9 signaling. *Infection and immunity*, *80*(3), 1274–1279. <https://doi.org/10.1128/IAI.06033-11>

Hahn, W. O., Butler, N. S., Lindner, S. E., Akilesh, H. M., Sather, D. N., Kappe, S. H., Hamerman, J. A., Gale, M., Jr, Liles, W. C., & Pepper, M. (2018). cGAS-mediated control of blood-stage malaria promotes *Plasmodium*-specific germinal center responses. *JCI insight*, *3*(2), e94142. <https://doi.org/10.1172/jci.insight.94142>

Haque, A., Best, S. E., Ammerdorffer, A., Desbarrieres, L., de Oca, M. M., Amante, F. H., de Labastida Rivera, F., Hertzog, P., Boyle, G. M., Hill, G. R., & Engwerda, C. R. (2011). Type I interferons suppress CD4⁺ T-cell-dependent parasite control during blood-stage *Plasmodium* infection. *European journal of immunology*, *41*(9), 2688–2698. <https://doi.org/10.1002/eji.201141539>

Hernandez-Valladares, M., Naessens, J., Musoke, A. J., Sekikawa, K., Rihet, P., Ole-Moiyoi, O. K., Busher, P., & Iraqi, F. A. (2006). Pathology of Tnf-deficient mice infected with *Plasmodium chabaudi adami* 408XZ. *Experimental parasitology*, *114*(4), 271–278. <https://doi.org/10.1016/j.exppara.2006.04.003>

Hou, N., Zou, Y., Piao, X., Liu, S., Wang, L., Li, S., & Chen, Q. (2016). T-cell immunoglobulin- and mucin-domain-containing molecule 3 signaling blockade improves cell-mediated immunity against malaria. *The Journal of infectious diseases*, *214*(10), 1547–1556. <https://doi.org/10.1093/infdis/jiw428>

Ing, R., & Stevenson, M. M. (2009). Dendritic cell and NK cell reciprocal cross talk promotes gamma interferon-dependent immunity to blood-stage *Plasmodium chabaudi* AS infection in mice. *Infection and immunity*, *77*(2), 770–782. <https://doi.org/10.1128/IAI.00994-08>

Ing, R., Gros, P., & Stevenson, M. M. (2005). Interleukin-15 enhances innate and adaptive immune responses to blood-stage malaria infection in mice. *Infection and immunity*, *73*(5), 3172–3177. <https://doi.org/10.1128/IAI.73.5.3172-3177.2005>

Jacobs, P., Radzioch, D., & Stevenson, M. M. (1995). Nitric oxide expression in the spleen, but not in the liver, correlates with resistance to blood-stage malaria in mice. *Journal of immunology (Baltimore, Md.: 1950)*, *155*(11), 5306–5313.

James, K. R., Soon, M. S. F., Sebina, I., Fernandez-Ruiz, D., Davey, G., Liligeto, U. N., et al. (2018). IFN Regulatory Factor 3 balances Th1 and T Follicular helper immunity during nonlethal blood-stage *Plasmodium* infection. *Journal of immunology (Baltimore, Md. : 1950)*, *200*(4), 1443–1456. <https://doi.org/10.4049/jimmunol.1700782>

Kim, C. C., Nelson, C. S., Wilson, E. B., Hou, B., DeFranco, A. L., & DeRisi, J. L. (2012). Splenic red pulp macrophages produce type I interferons as early sentinels of malaria infection but are dispensable for control. *PloS one*, *7*(10), e48126. <https://doi.org/10.1371/journal.pone.0048126>

Kobayashi, F., Ishida, H., Matsui, T., & Tsuji, M. (2000). Effects of in vivo administration of anti-IL-10 or anti-IFN-gamma monoclonal antibody on the host defense mechanism against *Plasmodium yoelii yoelii* infection. *The Journal of veterinary medical science*, *62*(6), 583–587. <https://doi.org/10.1292/jvms.62.583>

Maeno, Y., Nakazawa, S., Yamamoto, N., Shinzato, M., Nagashima, S., Tanaka, K., Sasaki, J., Rittling, S. R., Denhardt, D. T., Uede, T., & Taniguchi, K. (2006). Osteopontin participates in Th1-mediated host resistance against nonlethal malaria parasite *Plasmodium chabaudi* *chabaudi* infection in mice. *Infection and immunity*, *74*(4), 2423–2427. <https://doi.org/10.1128/IAI.74.4.2423-2427.2006>

Maglinao, M., Klopfleisch, R., Seeberger, P. H., & Lepenies, B. (2013). The C-type lectin receptor DCIR is crucial for the development of experimental cerebral malaria. *Journal of immunology (Baltimore, Md. : 1950)*, *191*(5), 2551–2559. <https://doi.org/10.4049/jimmunol.1203451>

Mannoor, M. K., Weerasinghe, A., Halder, R. C., Reza, S., Morshed, M., Ariyasinghe, A., Watanabe, H., Sekikawa, H., & Abo, T. (2001). Resistance to malarial infection is achieved by the cooperation of NK1.1(+) and NK1.1(-) subsets of intermediate TCR cells which are constituents of innate immunity. *Cellular immunology*, *211*(2), 96–104. <https://doi.org/10.1006/cimm.2001.1833>

Mastelic, B., do Rosario, A. P., Veldhoen, M., Renauld, J. C., Jarra, W., Sponaas, A. M., Roetynck, S., Stockinger, B., & Langhorne, J. (2012). IL-22 protects against liver pathology and lethality of an experimental blood-stage malaria infection. *Frontiers in immunology*, *3*, 85. <https://doi.org/10.3389/fimmu.2012.00085>

Muxel, S. M., Freitas do Rosário, A. P., Zago, C. A., Castillo-Méndez, S. I., Sardinha, L. R., Rodriguez-Málaga, S. M., Câmara, N. O., Álvarez, J. M., & Lima, M. R. (2011). The spleen CD4+ T cell response to blood-stage *Plasmodium chabaudi* malaria develops in two phases characterized by different properties. *PloS one*, *6*(7), e22434. <https://doi.org/10.1371/journal.pone.0022434>

Patel, S. N., Lu, Z., Ayi, K., Serghides, L., Gowda, D. C., & Kain, K. C. (2007). Disruption of CD36 impairs cytokine response to *Plasmodium falciparum* glycosylphosphatidylinositol and confers susceptibility to severe and fatal malaria in vivo. *Journal of immunology (Baltimore, Md. : 1950)*, *178*(6), 3954–3961. <https://doi.org/10.4049/jimmunol.178.6.3954>

Pattaradilokrat, S., Li, J., Wu, J., Qi, Y., Eastman, R. T., Zilversmit, M., Nair, S. C., Huaman, M. C., Quinones, M., Jiang, H., Li, N., Zhu, J., Zhao, K., Kaneko, O., Long, C. A., & Su, X. Z. (2014). Plasmodium genetic loci linked to host cytokine and chemokine responses. *Genes and immunity*, *15*(3), 145–152. <https://doi.org/10.1038/gene.2013.74>

Playfair JH. Lethal *Plasmodium yoelii* malaria: the role of macrophages in normal and immunized mice. Bulletin of the World Health Organization. 1979 ;57 Suppl 1:245-246.

Pung, O., & Katz, F. F. (1984). Primary lethal malaria (*Plasmodium chabaudi*) in hereditarily asplenic, splenectomized, and intact mice. *The Journal of parasitology*, *70*(2), 305–306.

Roland, J., Soulard, V., Sellier, C., Drapier, A. M., Di Santo, J. P., Cazenave, P. A., & Pied, S. (2006). NK cell responses to *Plasmodium* infection and control of intrahepatic parasite development. *Journal of immunology (Baltimore, Md. : 1950)*, *177*(2), 1229–1239. <https://doi.org/10.4049/jimmunol.177.2.1229>

Rudin, W., Eugster, H. P., Bordmann, G., Bonato, J., Müller, M., Yamage, M., & Ryffel, B. (1997). Resistance to cerebral malaria in tumor necrosis factor-alpha/beta-deficient mice is associated with a reduction of intercellular adhesion molecule-1 up-regulation and T helper type 1 response. *The American journal of pathology*, *150*(1), 257–266.

Rummel, T., Batchelder, J., Flaherty, P., LaFleur, G., Nanavati, P., Burns, J. M., & Weidanz, W. P. (2004). CD28 costimulation is required for the expression of T-cell-dependent cell-mediated immunity against blood-stage *Plasmodium chabaudi* malaria parasites. *Infection and immunity*, *72*(10), 5768–5774. <https://doi.org/10.1128/IAI.72.10.5768-5774.2004>

Sebina, I., James, K. R., Soon, M. S., Fogg, L. G., Best, S. E., Labastida Rivera, F., Montes de Oca, M., Amante, F. H., Thomas, B. S., Beattie, L., Souza-Fonseca-Guimaraes, F., Smyth, M. J., Hertzog, P. J., Hill, G. R., Hutloff, A., Engwerda, C. R., & Haque, A. (2016). IFNAR1-Signalling obstructs ICOS-mediated humoral immunity during non-lethal blood-stage *Plasmodium* infection. *PLoS pathogens*, *12*(11), e1005999. <https://doi.org/10.1371/journal.ppat.1005999>

Sellau, J., Alvarado, C. F., Hoenow, S., Mackroth, M. S., Kleinschmidt, D., Huber, S., & Jacobs, T. (2016). IL-22 dampens the T cell response in experimental malaria. *Scientific reports*, *6*, 28058. <https://doi.org/10.1038/srep28058>

Serghides, L., Patel, S. N., Ayi, K., Lu, Z., Gowda, D. C., Liles, W. C., & Kain, K. C. (2009). Rosiglitazone modulates the innate immune response to *Plasmodium falciparum* infection and improves outcome in experimental cerebral malaria. *The Journal of infectious diseases*, *199*(10), 1536–1545. <https://doi.org/10.1086/598222>

Singh, R. P., Kashiwamura, S., Rao, P., Okamura, H., Mukherjee, A., & Chauhan, V. S. (2002). The role of IL-18 in blood-stage immunity against murine malaria *Plasmodium yoelii* 265 and *Plasmodium berghei* ANKA. *Journal of immunology (Baltimore, Md. : 1950)*, *168*(9), 4674–4681. <https://doi.org/10.4049/jimmunol.168.9.4674>

Spaulding, E., Fooksman, D., Moore, J. M., Saidi, A., Feintuch, C. M., Reizis, B., Chorro, L., Daily, J., & Lauvau, G. (2016). STING-Licensed macrophages prime type I IFN production by plasmacytoid dendritic cells in the bone marrow during severe *Plasmodium yoelii* malaria. *PLoS pathogens*, *12*(10), e1005975. <https://doi.org/10.1371/journal.ppat.1005975>

Stevenson, M. M., & Ghadirian, E. (1989). Human recombinant tumor necrosis factor alpha protects susceptible A/J mice against lethal *Plasmodium chabaudi* AS infection. *Infection and immunity*, *57*(12), 3936–3939. <https://doi.org/10.1128/iai.57.12.3936-3939.1989>

Stevenson, M. M., Ghadirian, E., Phillips, N. C., Rae, D., & Podoba, J. E. (1989). Role of mononuclear phagocytes in elimination of *Plasmodium chabaudi* AS infection. *Parasite immunology*, *11*(5), 529–544. <https://doi.org/10.1111/j.1365-3024.1989.tb00687.x>

Stevenson, M. M., Tam, M. F., & Nowotarski, M. (1990). Role of interferon-gamma and tumor necrosis factor in host resistance to *Plasmodium chabaudi* AS. *Immunology letters*, *25*(1-3), 115–121. <https://doi.org/10.1016/0165-2478(90)90101-u>

Stevenson, M. M., Tam, M. F., & Rae, D. (1990). Dependence on cell-mediated mechanisms for the appearance of crisis forms during *Plasmodium chabaudi* AS infection in C57BL/6 mice. *Microbial pathogenesis*, *9*(5), 303–314. <https://doi.org/10.1016/0882-4010(90)90065-x>

Stevenson, M. M., Tam, M. F., Wolf, S. F., & Sher, A. (1995). IL-12-induced protection against blood-stage *Plasmodium chabaudi* AS requires IFN-gamma and TNF-alpha and occurs via a nitric oxide-dependent mechanism. *Journal of immunology (Baltimore, Md. : 1950)*, *155*(5), 2545–2556.

Su, Z., Fortin, A., Gros, P., & Stevenson, M. M. (2002). Opsonin-independent phagocytosis: an effector mechanism against acute blood-stage *Plasmodium chabaudi AS* infection. *The Journal of infectious diseases*, *186*(9), 1321–1329. <https://doi.org/10.1086/344576>

Süss, G., Eichmann, K., Kury, E., Linke, A., & Langhorne, J. (1988). Roles of CD4- and CD8-bearing T lymphocytes in the immune response to the erythrocytic stages of *Plasmodium chabaudi.* *Infection and immunity*, *56*(12), 3081–3088. <https://doi.org/10.1128/iai.56.12.3081-3088.1988>

Tamura, T., Akbari, M., Kimura, K., Kimura, D., & Yui, K. (2014). Flt3 ligand treatment modulates parasitemia during infection with rodent malaria parasites via MyD88- and IFN-γ-dependent mechanisms. *Parasite immunology*, *36*(2), 87–99. <https://doi.org/10.1111/pim.12085>

Tamura, T., Kimura, K., Yuda, M., & Yui, K. (2011). Prevention of experimental cerebral malaria by Flt3 ligand during infection with *Plasmodium berghei* ANKA. *Infection and immunity*, *79*(10), 3947–3956. <https://doi.org/10.1128/IAI.01337-10>

Taniguchi, T., Tachikawa, S., Kanda, Y., Kawamura, T., Tomiyama-Miyaji, C., Li, C., Watanabe, H., Sekikawa, H., & Abo, T. (2007). Malaria protection in beta 2-microglobulin-deficient mice lacking major histocompatibility complex class I antigens: essential role of innate immunity, including gammadelta T cells. *Immunology*, *122*(4), 514–521. <https://doi.org/10.1111/j.1365-2567.2007.02661.x>

Taverne, J., Sheikh, N., de Souza, J. B., Playfair, J. H., Probert, L., & Kollias, G. (1994). Anaemia and resistance to malaria in transgenic mice expressing human tumour necrosis factor. *Immunology*, *82*(3), 397–403.

Taverne, J., Tavernier, J., Fiers, W., & Playfair, J. H. (1987). Recombinant tumour necrosis factor inhibits malaria parasites in vivo but not in vitro. *Clinical and experimental immunology*, *67*(1), 1–4.

Theeß, W., Sellau, J., Steeg, C., Klinke, A., Baldus, S., Cramer, J. P., & Jacobs, T. (2016). Myeloperoxidase attenuates pathogen clearance during *Plasmodium yoelii* nonlethal Infection. *Infection and immunity*, *85*(1), e00475-16. <https://doi.org/10.1128/IAI.00475-16>

Thylur, R. P., Wu, X., Gowda, N. M., Punnath, K., Neelgund, S. E., Febbraio, M., & Gowda, D. C. (2017). CD36 receptor regulates malaria-induced immune responses primarily at early blood stage infection contributing to parasitemia control and resistance to mortality. *The Journal of biological chemistry*, *292*(22), 9394–9408. <https://doi.org/10.1074/jbc.M117.781294>

Tsutsui, N., & Kamiyama, T. (1998). Suppression of in vitro IFN-gamma production by spleen cells of *Plasmodium chabaudi*-infected C57BL/10 mice exposed to dexamethasone at a low dose. *International journal of immunopharmacology*, *20*(4-5), 141–152. <https://doi.org/10.1016/s0192-0561(98)00019-8>

Tsutsui, N., & Kamiyama, T. (1999). Transforming growth factor beta-induced failure of resistance to infection with blood-stage *Plasmodium chabaudi* in mice. *Infection and immunity*, *67*(5), 2306–2311. <https://doi.org/10.1128/IAI.67.5.2306-2311.1999>

Van der Heyde, H. C., Gu, Y., Zhang, Q., Sun, G., & Grisham, M. B. (2000). Nitric oxide is neither necessary nor sufficient for resolution of Plasmodium chabaudi malaria in mice. *Journal of immunology (Baltimore, Md. : 1950)*, *165*(6), 3317–3323. <https://doi.org/10.4049/jimmunol.165.6.3317>

Vigário, A. M., Belnoue, E., Grüner, A. C., Mauduit, M., Kayibanda, M., Deschemin, J. C., Marussig, M., Snounou, G., Mazier, D., Gresser, I., & Rénia, L. (2007). Recombinant human IFN-alpha inhibits cerebral malaria and reduces parasite burden in mice. *Journal of immunology (Baltimore, Md. : 1950)*, *178*(10), 6416–6425. <https://doi.org/10.4049/jimmunol.178.10.6416>

Villegas-Mendez, A., Inkson, C. A., Shaw, T. N., Strangward, P., & Couper, K. N. (2016). Long-Lived CD4+IFN-γ+ T Cells rather than Short-Lived CD4+IFN-γ+IL-10+ T Cells Initiate Rapid IL-10 Production To Suppress Anamnestic T Cell Responses during Secondary Malaria Infection. *Journal of immunology (Baltimore, Md. : 1950)*, *197*(8), 3152–3164. <https://doi.org/10.4049/jimmunol.1600968>

Voisine, C., Mastelic, B., Sponaas, A. M., & Langhorne, J. (2010). Classical CD11c+ dendritic cells, not plasmacytoid dendritic cells, induce T cell responses to Plasmodium chabaudi malaria. *International journal for parasitology*, *40*(6), 711–719. <https://doi.org/10.1016/j.ijpara.2009.11.005>

Weidanz, W. P., LaFleur, G., Brown, A., Burns, J. M., Jr, Gramaglia, I., & van der Heyde, H. C. (2010). Gammadelta T cells but not NK cells are essential for cell-mediated immunity against Plasmodium chabaudi malaria. *Infection and immunity*, *78*(10), 4331–4340. <https://doi.org/10.1128/IAI.00539-10>

Weidanz, W. P., Lafleur, G., Kita-Yarbro, A., Nelson, K., & Burns, J. M., Jr (2011). Signalling through the IL-2 receptor γ(c) peptide (CD132) is essential for the expression of immunity to *Plasmodium chabaudi adami blood*-stage malaria. *Parasite immunology*, *33*(9), 512–516. <https://doi.org/10.1111/j.1365-3024.2011.01298.x>

Wikenheiser, D. J., Brown, S. L., Lee, J., & Stumhofer, J. S. (2018). NK1.1 Expression Defines a Population of CD4^+^ Effector T Cells Displaying Th1 and Tfh Cell Properties That Support Early Antibody Production During *Plasmodium yoelii* Infection. *Frontiers in immunology*, *9*, 2277. <https://doi.org/10.3389/fimmu.2018.02277>

Wu, J., Xia, L., Yao, X., Yu, X., Tumas, K. C., Sun, W., Cheng, Y., He, X., Peng, Y. C., Singh, B. K., Zhang, C., Qi, C. F., Bolland, S., Best, S. M., Gowda, C., Huang, R., Myers, T. G., Long, C. A., Wang, R. F., & Su, X. Z. (2020). The E3 ubiquitin ligase MARCH1 regulates antimalaria immunity through interferon signaling and T cell activation. *Proceedings of the National Academy of Sciences of the United States of America*, *117*(28), 16567–16578. <https://doi.org/10.1073/pnas.2004332117>

Wu, X., Dayanand, K. K., Thylur, R. P., Norbury, C. C., & Gowda, D. C. (2017). Small molecule-based inhibition of MEK1/2 proteins dampens inflammatory responses to malaria, reduces parasite load, and mitigates pathogenic outcomes. *The Journal of biological chemistry*, *292*(33), 13615–13634. <https://doi.org/10.1074/jbc.M116.770313>

Wu, X., Thylur, R. P., Dayanand, K. K., Punnath, K., Norbury, C. C., & Gowda, D. C. (2021). IL-4 Treatment Mitigates Experimental Cerebral Malaria by Reducing Parasitemia, Dampening Inflammation, and Lessening the Cytotoxicity of T Cells. *Journal of immunology (Baltimore, Md. : 1950)*, *206*(1), 118–131. <https://doi.org/10.4049/jimmunol.2000779>

Wunderlich, F., Dkhil, M. A., Mehnert, L. I., Braun, J. V., El-Khadragy, M., Borsch, E., Hermsen, D., Benten, W. P., Pfeffer, K., Mossmann, H., & Krücken, J. (2005). Testosterone responsiveness of spleen and liver in female lymphotoxin beta receptor-deficient mice resistant to blood-stage malaria. *Microbes and infection*, *7*(3), 399–409. <https://doi.org/10.1016/j.micinf.2004.11.016>

Yao, X., Wu, J., Lin, M., Sun, W., He, X., Gowda, C., Bolland, S., Long, C. A., Wang, R., & Su, X. Z. (2016). Increased CD40 Expression Enhances Early STING-Mediated Type I Interferon Response and Host Survival in a Rodent Malaria Model. *PLoS pathogens*, *12*(10), e1005930. <https://doi.org/10.1371/journal.ppat.1005930>

Zander, R. A., Guthmiller, J. J., Graham, A. C., Pope, R. L., Burke, B. E., Carr, D. J., & Butler, N. S. (2016). Type I Interferons Induce T Regulatory 1 Responses and Restrict Humoral Immunity during Experimental Malaria. *PLoS pathogens*, *12*(10), e1005945. <https://doi.org/10.1371/journal.ppat.1005945>

Zhang, Y., Zhu, X., Feng, Y., Pang, W., Qi, Z., Cui, L., & Cao, Y. (2016). TLR4 and TLR9 signals stimulate protective immunity against blood-stage Plasmodium yoelii infection in mice. *Experimental parasitology*, *170*, 73–81. <https://doi.org/10.1016/j.exppara.2016.09.003>
